# Transcriptomic profiling reveals neurophysiological gene candidates underlying vocal evolution in African clawed frogs

**DOI:** 10.64898/2026.01.24.701475

**Authors:** Charlotte Barkan, Lucas Binder, Brett Davis, Lucia Carbone, Erik Zornik

## Abstract

Neurophysiologists have discovered many mechanisms underlying the production of animal behaviors in specific species; these involve a collection of neuromuscular systems, neuronal membrane and neural network properties, as well as the hormones and neuromodulators known to modify them. However, the mechanistic basis of behavioral evolution is less well-studied, and causal links between differences in gene expression, cellular mechanisms and species-typical behaviors are rare. Vertebrate vocal behaviors are an excellent system for studying the evolution of behaviors because they are ancient, diverse and readily quantifiable. *Xenopus* frogs are particularly well-suited to the study of vocal evolution due to the temporal diversity of male advertisement calls between closely related species and the well-described vocal pattern generating circuitry. Here we focus on two species, *X. laevis* and *X. petersii*, that diverged 8.5 million years ago and produce advertisement calls with distinct timing. To begin bridging the gap between behavioral and mechanistic diversity in *Xenopus* vocal behaviors, we performed RNA sequencing of the parabrachial nucleus, a vocal premotor hindbrain area known to encode species-typical temporal patterns in *X. laevis* and *X. petersii*. We identified hundreds of differentially expressed genes between the two species, including many genes related to hormone signaling, neuromodulation, neuronal and synaptic functions, ion channels and neurotransmitter receptors. We explore several testable hypotheses emerging from these results that may explain mechanisms by which candidate genes and gene families may contribute to vocal pattern differences between *X. laevis* and *X. petersii*.

## INTRODUCTION

Vertebrate behaviors are generated by coordinated activation of muscles controlled by neuronal circuits in the spinal cord and brain. The diversity of animal behaviors that exist are the result of evolution of these behavioral effectors including muscles, motor neurons and pattern-generating circuits. Consequently, evolutionary changes in the properties of these circuits can give rise to differences in behavior. Neuronal circuits can evolve by changes in A) connectivity, synaptic strength and neuron number (1–3), B) neuromodulation (4,5), and C) neuronal physiology (6–7) see 8 for detailed review). Pinpointing the genetic basis of evolutionary adaptations and linking them to specific neural circuit properties is a major goal of evolutionary neurobiology.

Acoustic communication is observed across all major clades of vertebrates (fish, amphibians, reptiles, birds and mammals) and plays essential roles in coordinating intraspecific interactions including courtship, social structures, territoriality and parental care (9–16). Acoustic signals across animals are highly diverse and quantifiable, making them an effective system for studying behavioral evolution. While several recent studies have begun to identify neuronal (7) and genetic (6) mechanisms underlying evolution of communication signals, it remains rare to be able to link the two together (but see 17).

Vocalizations of *Xenopus* frogs are a powerful model in which to identify evolutionary links between genes, neuronal circuits, and behavior. The *Xenopus* phylogeny contains ∼30 described species distributed across sub-Saharan Africa (18,19). Males of each species produce a unique advertisement call defined by its spectrotemporal properties that attract females of their own species during courtship (20). The temporal aspects of this call are primarily controlled by a hindbrain central pattern generator (7,21–23). Premotor neurons in the *Xenopus* hindbrain parabrachial nucleus (PB) project directly to the vocal motor nucleus; motor neurons in turn send their axons via nerve IX-X to a pair of laryngeal muscles responsible for activating sound production (24,25).

Previous work pinpointed premotor PB as a site of species differences in generating advertisement call duration and period in *X. laevis* and and *X. petersii* (7,21–23). *X. laevis* produces an advertisement call with a long duration and period compared with a shorter duration and period in *X. petersii* (Fig. 1). When PB is physically isolated from other vocal areas, these species-specific rhythms persist autonomously in the local circuit (23). Furthermore, synaptically isolated PB neurons can generate membrane potential oscillations with species-specific temporal patterns (7). This evidence suggests that there are intrinsic differences in vocal PB neuron expression and/or function of functionally important proteins.

**Figure 1.**
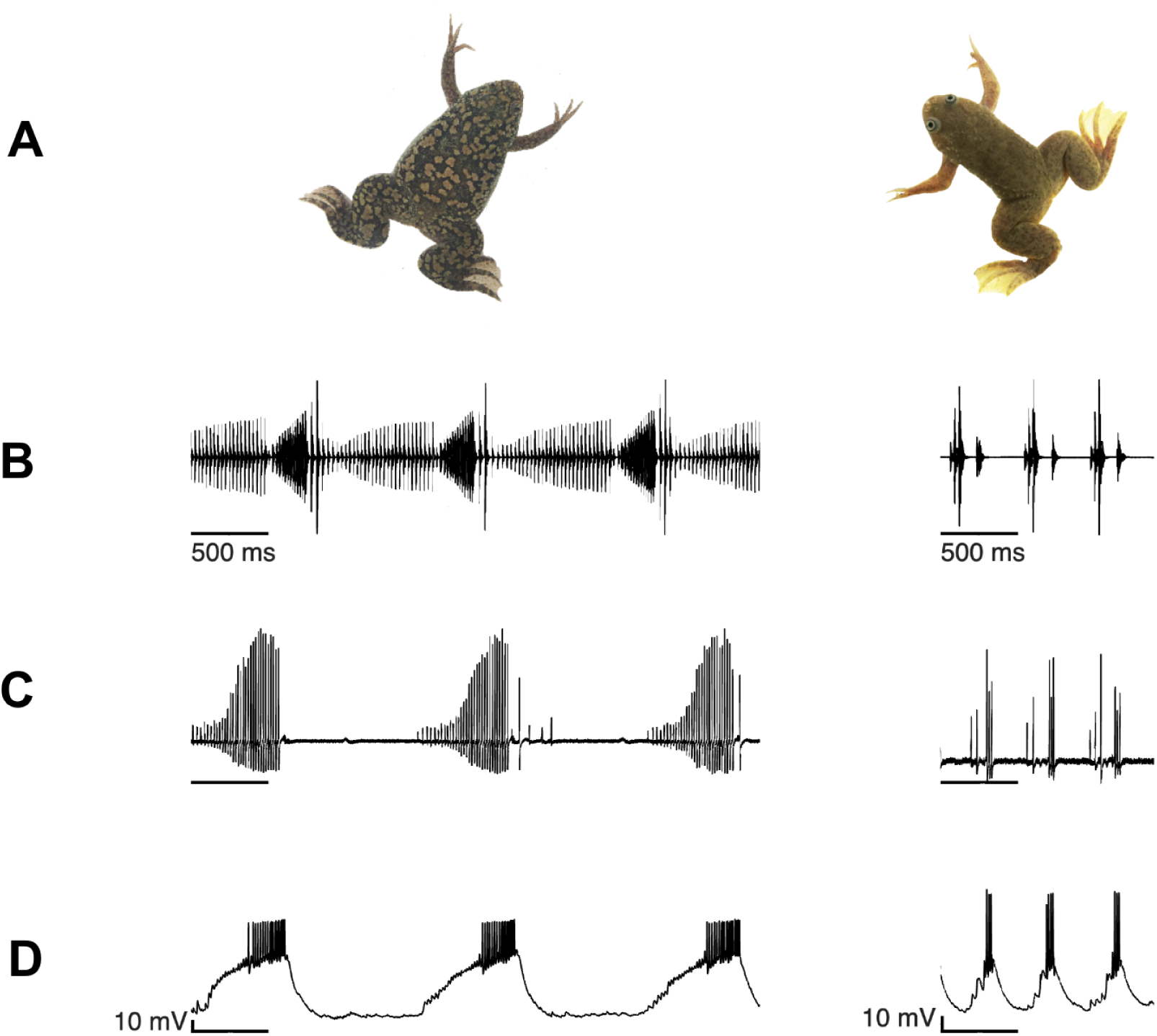
Xenopus advertisement calls vary in temporal properties across species. (A) Male *X. laevis* (right) and male X. petersii (left) frog. (B) In vivo advertisement call of X. laevis (right) and X. petersii (left). In both species advertisement calls consist of a slow trill (∼ 30 Hz sound pulses) followed by a fast trill (∼ 60 Hz), however the trill duration and call period of *X. laevis* is significantly longer than duration and period of X*. petersii* calls. (C) Ex vivo fictive advertisement calls produced by isolated X. laevis (right) and X. petersii (left) brains. Trill rates, durations and call period are similar in ex vivo preparation to those in in vivo song. (D) Premotor parabrachial nucleus (PB) fast trill neurons during fictive call in *X. laevis* (right) and *X. petersii* (left). PBx neurons depolarize and spike during fast trill.

To determine the degree to which differences in expression of ion channels and neurotransmitter receptors contribute to functional differences in the premotor vocal circuit, we used RNA sequencing to compare gene expression patterns in PB of *X. laevis* and *X. petersii* males. We hypothesized that species-typical vocal patterns would be reflected in the kinetics of the ion channels and neurotransmitter receptors encoded by differentially expressed genes. In addition, because *Xenopus* vocal behaviors are hormone-dependent, sex-biased, and temporally dynamic, we explore the potential impact of species-biased expression of hormone and neuromodulatory pathways on vocal evolution.

## METHODS

### Animals

Sexually mature, age-matched *X. laevis* (n=10; weight: 49.8 ± 4.4 g, length: 7.8 ± 0.3 cm and *X. petersii* (n=10; weight: 13.0 ± 1.0 g; length: 4.9 ± 0.2) male frogs were used for all experiments. *X. laevis* and *X. petersii* were purchased from Xenopus Express. Animals were group housed with a recirculating water system in PETG aquaria (Aquaneering) on a 12 h light/12 h dark schedule at 20°C and fed twice weekly. All animal care and experimental procedures conformed to guidelines of the National Institutes of Health and were approved by Reed College’s Institutional Animal Care and Use Committee (protocol no. 022017).

### Tissue collection and RNA extraction

Frogs were deeply anesthetized by injection of 1.3% tricaine methanesulfonate (MS-222; Sigma-Aldrich; X. laevis: 500 µl, X. petersii: 200 µl) into the dorsal lymph sac. Brains were rapidly dissected from the skull in ice-cold saline containing the following (in mM): 96 NaCl, 20 NaHCO_3_, 2 CaCl_2_, 2 KCl, 0.5 MgCl_2_, 10 HEPES, and 11 glucose, pH 7.8, oxygenated with 99% O_2_ (1% CO_2_) and stored overnight in RNAlater (Invitrogen) at 4°C.

The following day brains were vibratomed (Leica VT1000S) while submerged in RNAlater. A 400 µM horizontal section containing PB, beginning just ventral to the cerebellum, was collected and briefly stored on ice in RNAlater and then pinned in a silicone elastomer-lined recording dish (Sylgard; Dow Corning) submerged in RNAlater. A 22 gauge blunt syringe tip (Component Supply) was used to isolate and remove the region of the brain containing PB. The tissue punch was transferred to RNA extraction buffer (200 µL homogenization buffer with 4 µL thioglycerol, Maxwell 16 LEV simplyRNA Tissue Kit, Promega). RNA was extracted using the Maxwell 16 Instrument configured with the Maxwell 16 High Strength LEV Magnetic Rod and Plunger Bar Adaptor and then stored immediately at -80°C. Quality control was performed on the Bioanalyzer (Agilent) and RIN scores using the Total RNA Pico Chip. Punched brain sections were fixed in 4% paraformaldehyde, cryostat sectioned, stained with cresyl violet (0.1 %), and imaged to confirm punch site accuracy.

### Library preparation and sequencing

RNA-seq libraries were generated by the OHSU Massively Parallel Sequencing Shared Resources (MPSSR) using the Takara SmartSeq Plus kit and sequencing was performed on the Illumina NovaSeq 6000 (S4 flow cell).

### Data processing and differential expression analyses

All samples were aligned to the *Xenopus laevis* genome v10.1 (XenLae10) obtained from NCBI with the STAR aligner, along with NCBI RefSeq annotation. Alignment rates varied slightly between species, with *X. laevis* samples having around 91% alignment and *X. petersii* samples having around 85% alignment (Supplementary table 1).

We used the software edgeR (v3.28.0) for differential gene expression analysis (26). As input, we used a gene counts matrix with samples as columns, genes as rows (low count filtered genes), and the corresponding counts filling the cells of the table. Our low-count filtering criteria resulted in ∼26,000 genes being retained and considered for differential expression analysis. Our variable of interest was the species (*X. laevis* or *X. petersii*) with no covariates. Genes with an adjusted p-value < 0.05 (BH adjusted for multiple testing) and a log_2_ fold change (LFC) value > 1 were designated as significantly upregulated in *X. petersii*, and genes with an adjusted p-value < 0.05 and a log fold change value < -1 were designated as significantly upregulated in *X. laevis*.

### Gene-category and functional pathway analyses

Gene Ontology (GO) term and KEGG pathway enrichment analyses were performed using the clusterProfiler package (27) version 4.8.3 and the *Xenopus laevis* genome annotation org.Xl.eg.db version 3.17.0 (28) in R Studio. Principal component analysis between samples was performed using the scikit-learn Python package (29). In order to explore a comprehensive list of genes related to synaptic signaling (including ionotropic and metabotropic receptors, neuropeptides and protein hormones) we identified all genes in our dataset belonging to the “neuroactive ligand-receptor interaction” KEGG pathway. Genes encoding pore-forming and modulatory subunits in the voltage-gated ion channel family were extracted from the data set using the KEGG BRITE list for “Ligand-gated channels” and “Voltage-gated cation channels”. Genes encoding hormone receptors and steroid hormone biosynthesis were selected from the “neuroactive ligand-receptor interaction” and “Steroid hormone biosynthesis” KEGG pathways respectively.

## RESULTS

### Differential expression and enrichment analyses

Sequencing reads were mapped and filtered based on quality as described above (see Methods). Using EdgeR, a total of 6539 genes (24.6%) were found to be differentially expressed between species (padj < 0.05, Log_2_FC >= 1; Fig. 2). There were more down-regulated genes (3762; 57.5%) than up-regulated genes (2777; 42.5%) in *X. petersii* compared with *X. laevis* than expected by chance (Fisher’s exact test, p<0.00001). Principal component analysis (PCA) of the complete dataset separated the two species along the first principal component, which explained 39.37% of the variation in the data (Fig. 2).

**Figure 2.**
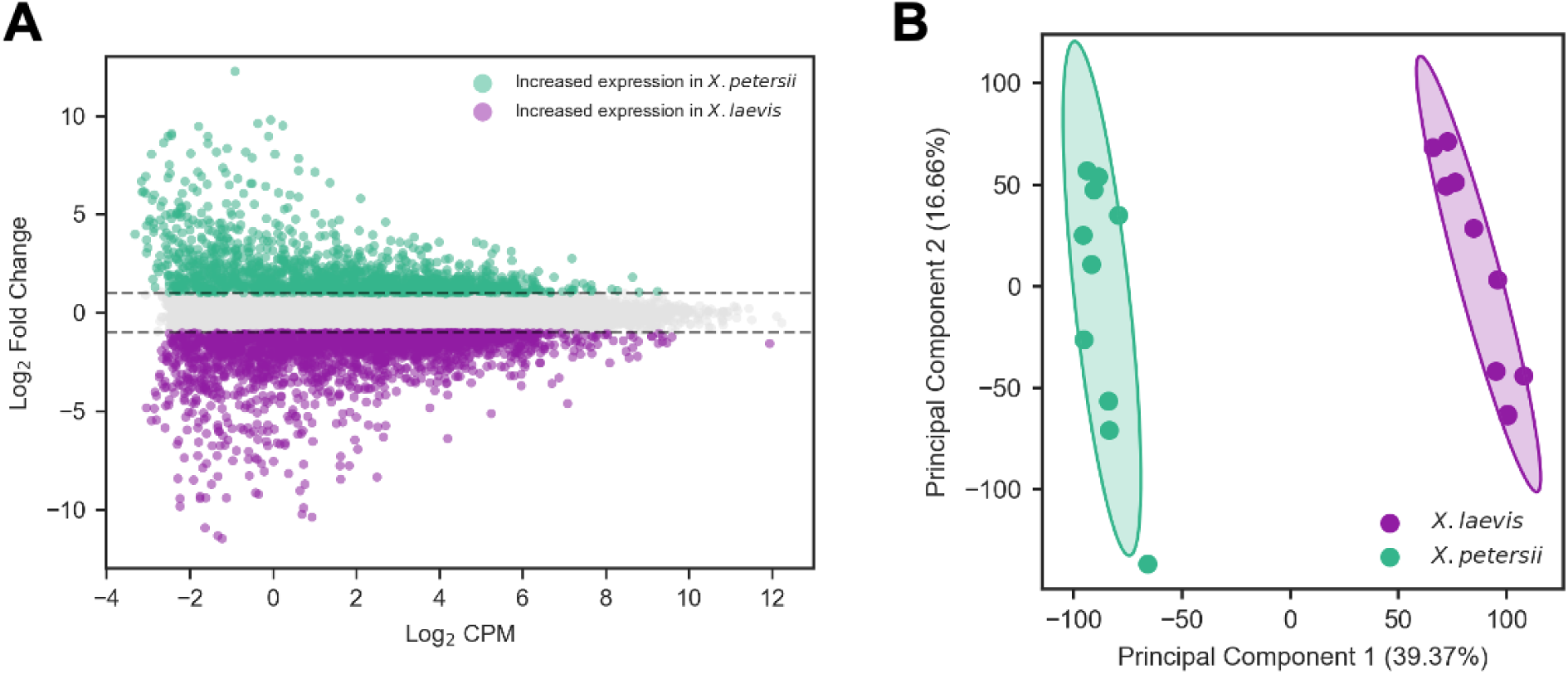
Transcriptomic differences between X. laevis and X. petersii. (A) Differential expression analysis identified 2777 upregulated genes in X. petersii (cyan) and 3762 genes upregulated in X. laevis (magenta). (B) Principal component analysis (with 95% confidence interval ellipses) of the complete normalized transcriptome (26576 genes) reveal strong separation of the two species along PC1, accounting for over 39% of the dataset variation.

We used GO term and KEGG pathway enrichment analyses to identify overrepresentation of gene categories and pathways among the 6539 differentially expressed genes (DEGs). Using a low stringency cutoff (adjusted p-value < 0.2), we identified 40 GO terms related to Molecular Function, 22 Biological Processes terms, and 4 terms related to Cellular Compartment (Supplemental Data Table 2.1-2.3); we also identified 22 overrepresented KEGG pathways (Supplemental Data Table 3).

### Hormone signaling

Vocal behaviors in *Xenopus* are known to be hormonally regulated (30). As expected, DEGs were enriched for multiple hormone-related GO terms, including “hormone activity”, “eicosanoid receptor activity”, and “monooxygenase activity” as well as KEGG pathways related to hormone metabolism including “steroid hormone biosynthesis” and “arachidonic acid metabolism” (containing genes involved in prostaglandin synthesis). Here we explore hormone-related gene candidates that may underlie behavioral differences between *X. laevis* and *X. petersii*.

Nuclear steroid receptor genes were expressed in PB of both species, including androgen, estrogen, progesterone, glucocorticoid and mineralocorticoid receptors (Fig. 3A). ER𝛼 (esr1.L) and progesterone receptor (pgr.L) transcripts were upregulated in *X. petersii*

**Figure 3.**
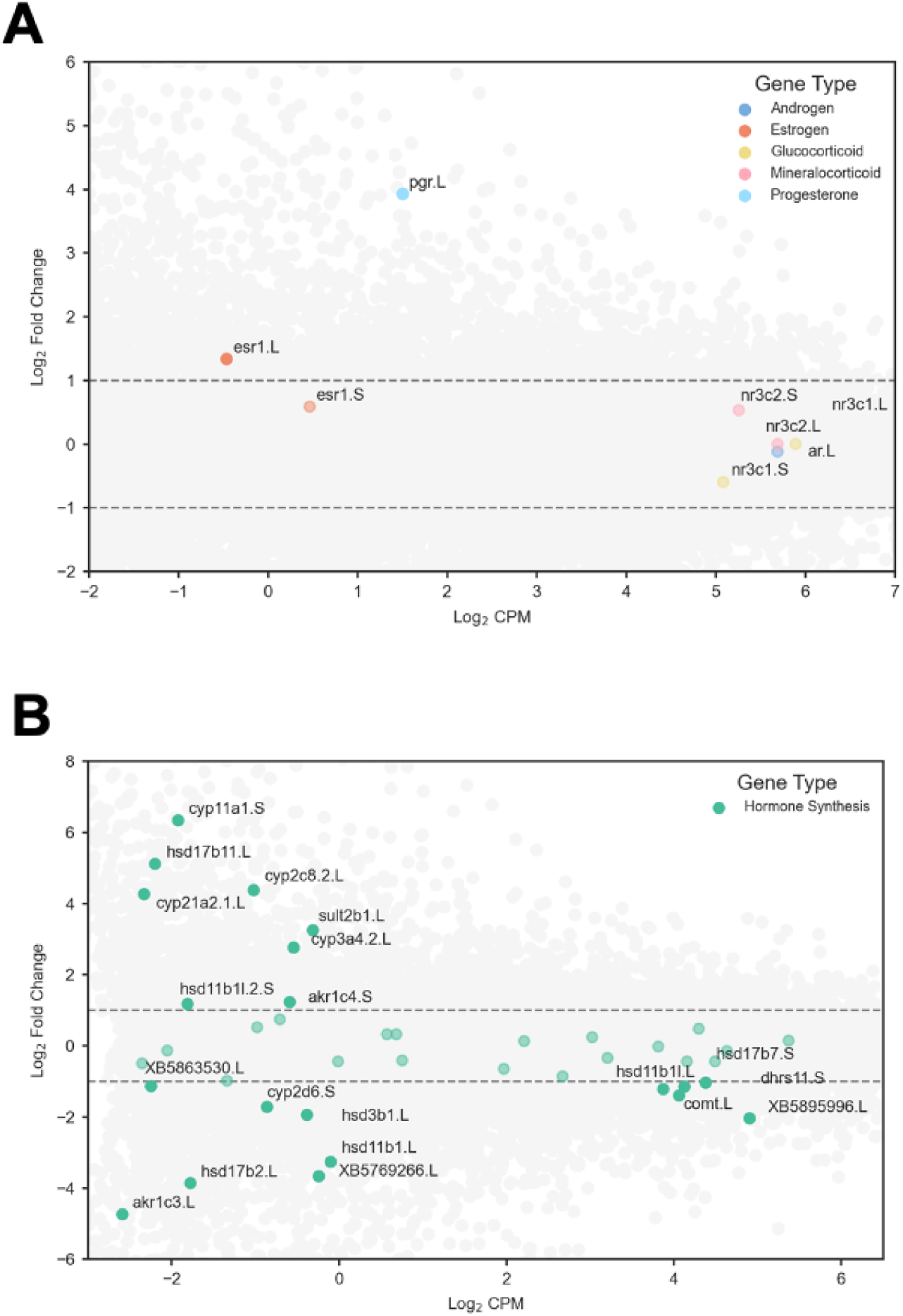
Hormone pathway gene expression. (A) All classes of nuclear steroid hormone receptors are expressed in PB of both *X. laevis* and *X. petersii*. The progesterone receptor and one of two estrogen receptor alpha transcripts were upregulated in *X. petersii*. (B) Genes encoding enzymes throughout the steroid synthesis pathways are expressed in PB; 50% of hormone synthesis genes were differentially expressed. In these and all subsequent MA plots, positive LFC values represent genes upregulated in *X. petersii*; negative LFC values represent genes upregulated in *X. laevis*.

Genes encoding steroid synthesis and metabolism enzymes were expressed (Fig. 3B). The gene encoding the cholesterol side-chain cleavage enzyme (cyp11a1.S)–the entry point for the entire steroid pathway–was highly upregulated in *X. petersii* compared with *X. laevis* (LFC 6.35). Additional differentially expressed genes in the steroid pathways included 3-𝛽-hydroxysteroid dehydrogenase (3𝛽-HSD; hsd3b1.L), multiple 17𝛽-HSD enzymes (hsd17b2.L, hsd17b6.L, hsd17b7.L, hsd17b7.S, and hsd17b11.L), and 21-hydroxylase (cyp21a2.1.L). Genes encoding other key steroid metabolic enzymes like aromatase (cyp19a1.L) and 5𝛼-reductase (srd5a1.S, srd5a3.L, and srd5a3.S) were present but not differentially expressed.

Multiple genes involved in prostaglandin metabolism were differentially expressed; including prostaglandin D2 synthase (ptgds.S), prostaglandin I synthase (ptgis.L), and prostaglandin reductase 1 (ptgr1.2.L; downregulated in *X. petersi)*; as well as the prostaglandin endoperoxide synthase 2 gene (ptgs2.L; upregulated in *X. petersii*).

Differentially expressed non-steroid hormone receptors included genes encoding four of the six prostaglandin E receptors (upregulated in *X. petersii*: ptegr1.L, ptger3.L, ptger4.L; downregulated in *X. petersii*: ptger2.S) and the prostaglandin F2𝛼 receptor (ptgfr.L; upregulated in *X. petersii*). The melatonin receptor (mtnr1c.S) was also upregulated in *X. petersii*.

### Neuropeptide and neuromodulator pathways

Multiple GO terms and KEGG pathways related to synaptic function and signaling were overexpressed in the differential expression analyses, including “ligand-gated monoatomic cation channel activity”, and “G protein-coupled receptor activity”, suggesting that changes in neuromodulatory pathways may contribute to vocal evolution in *Xenopus*.

Differentially expressed neuropeptide receptor genes (Fig. 4A) included those coding for vasopressin receptors (avpr2.L, avpr2.S; both downregulated), angiotensin receptors (agtr1.L, agtr2.L; both downregulated), and galanin receptors (galr1.S, upregulated; galr2.S downregulated). Several genes encoding neuropeptides and protein hormones were differentially expressed (Fig. 4B). Genes coding for galanin (gal.1.L) and prolactin-releasing peptide (prlh.L) were downregulated in *X. petersii;* genes encoding the peptides vasopressin (avp.L, avp.S), neuropeptide Y (npy.L, npy.S), and angiotensin (agt.L) were upregulated. Two hormone peptides that participate in the hypothalamic-pituitary-gonadal pathway–follicle stimulating hormone (fshb.S) and gonadotropin-releasing hormone (gnrh1.L)–were downregulated in *X. petersii*.

**Figure 4.**
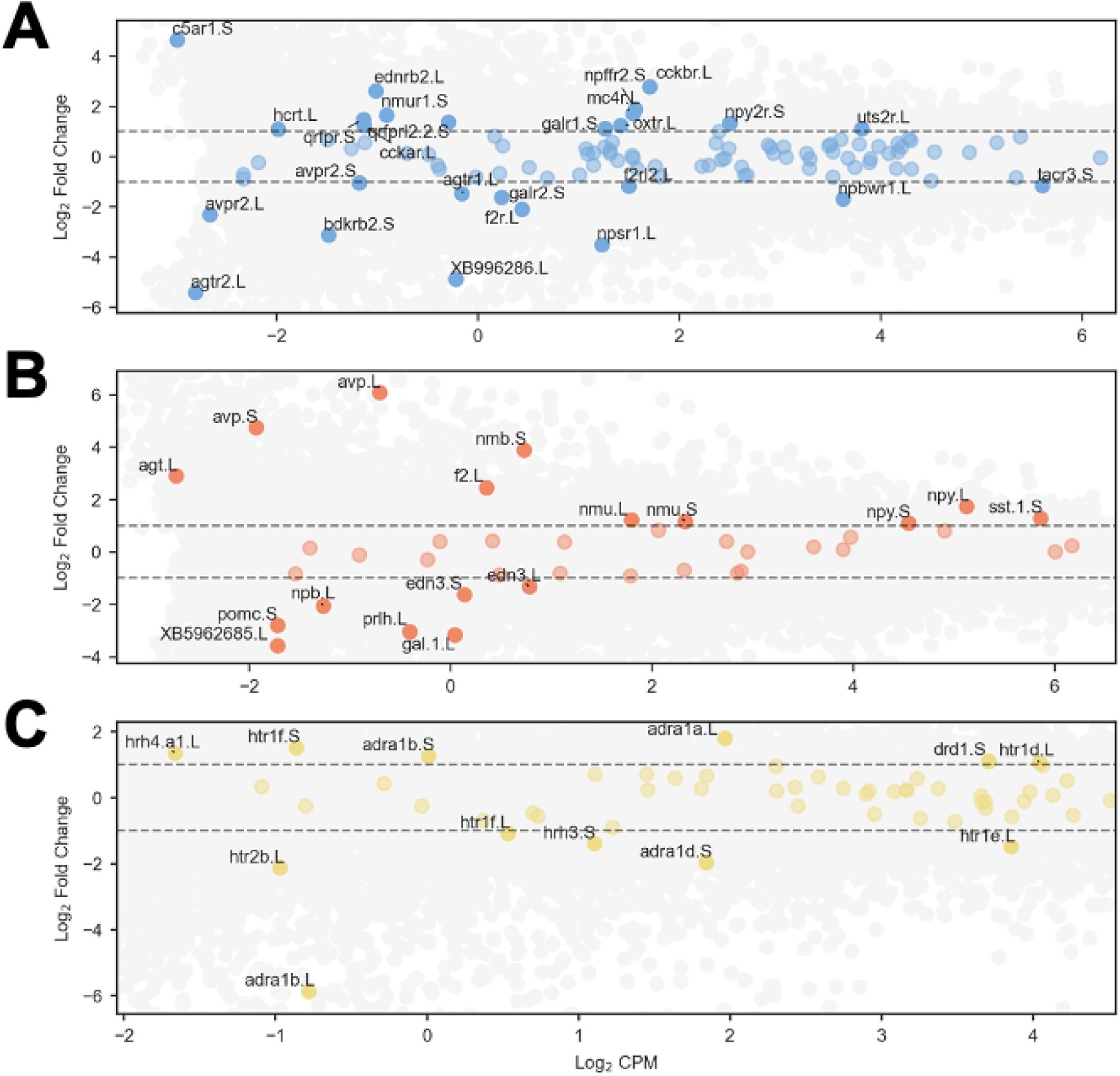
Neuropeptide and monoamine neuromodulator pathways. Many differentially expressed neuropeptide receptors (A) and neuropeptides (B) were identified in PB. Monoamine receptors (C) for serotonin, norepinephrine, and dopamine were differentially expressed between species.

Several genes encoding neuromodulatory amine receptors were expressed in the *Xenopus* vocal circuit (Fig. 4C). Six of 28 serotonin receptors and 1 of 9 dopamine receptors, were differentially expressed. 12 adrenergic receptors were identified, 4 of which were differentially expressed, while 2 of 6 histamine receptors were differentially expressed.

### Ion channels and neurotransmitter receptors

Temporal features of vocal rhythms depend on cell autonomous properties that differ between species (7); therefore we hypothesized that genes regulating membrane physiology and neural circuit properties are likely involved in vocal evolution. Consistent with this hypothesis, enriched GO terms included “channel activity”, “ligand-gated monoatomic cation channel activity”, and “passive transmembrane transporter activity”. To further explore the role of ion channels in vocal production and evolution across *X. laevis* and *X. petersii*, we identified 168 members of the voltage-gated (and structurally related) superfamily of genes as well as genes coding for regulatory subunits of the pore-forming channels (see Methods for details; Supplemental Data Table 3). Below we explore key gene candidates that may contribute to vocal neuron property differences between *X. laevis* and *X. petersii*.

We identified 95 genes encoding voltage-gated potassium channels and their regulatory subunits, 26 of which were differentially expressed (Fig. 5A). Over half of the 15 inwardly rectifying potassium channels were differentially expressed, 5 were downregulated (kcnj8.L, kcnj8.S, kcnj10.L, kcnj10.S, kcnj16.S) and 3 were upregulated (kcnj4.L, kcnj12.L, kcnj13.S) in *X. petersii.* Of the 16 tandem pore domain potassium leak channel genes, 5 were differentially expressed: 2 upregulated in *X. petersii* (kcnk13.L, kcnk1.S) and 3 downregulated in *X. petersii* (kcnk5.S, kcnk15.S, kcnk6.S). One small conductance calcium-activated potassium channel (SK) gene (kcnn3.S) was expressed in PB and was downregulated in *X. petersii*. While the one big conductance calcium-activated potassium channel (BK) gene expressed in PB (kcnma1.L) was not differentially expressed, 2 of the 5 genes coding for BK beta subunits were differentially expressed: kcnmb3.S was upregulated and kcnmb1.S was downregulated in *X. petersii*. Four of the eight genes Kv7 channels were differentially expressed: kcnq1.L and kcnq1.S were downregulated in X. petersii; kcnq3.L and kcnq3.S, which contribute to the M current, were upregulated in *X. petersii*. Other voltage-gated potassium channel superfamily genes were also differentially expressed including those that mediate A-type currents (kcnd1.S) and sodium-activated potassium channels (kcnt2l.L).

**Figure 5.**
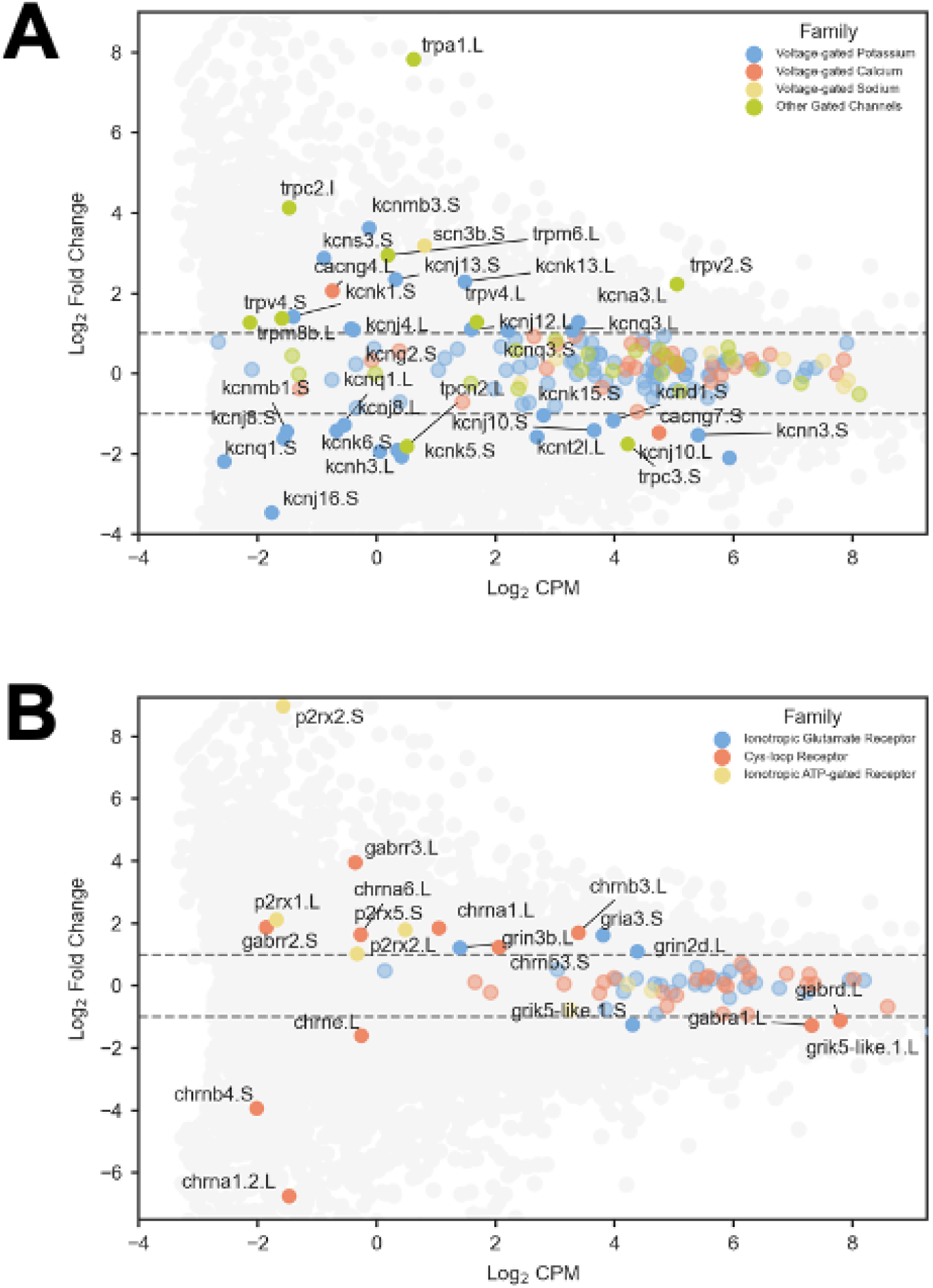
Ion channels and ionotropic neurotransmitter receptors. (A) We detected 168 genes encoding ion channels and modulatory subunits in the voltage-gated super family. Genes encoding voltage-gated potassium channel and transient receptor potential channel subunits were highly differentially expressed (27% and 36%, respectively); sodium (12.5%) calcium (6%) channel subunit genes showed the least differential expression. (B) Examples of differential expressed ionotropic receptor genes were identified for glutamatergic, GABAergic, cholinergic and purinergic receptors.

Only one gene coding for a voltage-gated sodium channel alpha subunit was expressed (and not differentially expressed): scn1a.S. Of the seven genes coding for sodium channel beta subunits, one was differentially expressed (scn3b.S was upregulated in *X. petersii*).

Of the 33 genes coding for voltage-gated calcium channel complexes, only two genes, both coding for gamma subunits, were differentially expressed: cacng7.S was downregulated, and cacng4.L was upregulated, in *X. petersii* compared to *X. laevis*. Three genes encoding pore-forming L-type channel subunits were present in the current dataset; although none reached the LFC threshold of the current study, all three genes exhibited modestly increased expression in *X. petersii*–Ca_V_1.2: cacna1c.L (LFC 0.48), cacna1c.S (LFC: 0.73); Ca_V_1.3: cacna1d.L (LFC: 0.73).

Both species showed high expression of the hyperpolarization-activated cation non-specific channel genes hcn1 (hcn1.L and hcn1.S), neither of which was differentially expressed. Likewise, none of the three observed cyclic nucleotide-gated ion channel genes (cnga3.L, cnga3.S, cngb1.S) was differentially expressed._Of the 22 genes encoding transient receptor potential (trp) channels, eight were differentially expressed, with seven upregulated, including trpa1.L, trpc2.L and trpm6.L, and one downregulated (trpc3.S) in *X. petersii*.

We identified 74 ionotropic transmitter receptor genes expressed in PB, 19 of which were differentially expressed (Fig. 5B). A total of 22 ionotropic glutamate receptor genes were identified, three of which were differentially expressed between the species; one AMPA receptor (gria3.S) and two NMDA receptor (grin2b.L and grin3b.L) were upregulated in *X. petersii*. None of the seven kainate receptors were differentially expressed. We identified 18 genes that encode nicotinic acetylcholine receptors (nAChR) complexes. Four genes were upregulated (chrna1.L, chrnb3.L, chrna6.L, chrnb3.S) while three were downregulated (chrne1.L, chrnb4.S, chrna1.2.L) in *X. petersii*. Among fast inhibitory receptors, 13 GABA-A genes were expressed, with one downregulated (gabra1.L) and two upregulated (gabrr3.L and gabrr2.S) in *X. petersii*. While we identified six glycine receptor genes, none of these was differentially expressed.

## DISCUSSION

### Overview

Behavioral differences can arise from evolutionary changes to the underlying neuronal circuits, however, few studies have identified differences between circuits that underlie species-specific behaviors (7,31). In the current study, we chose an unbiased RNA sequencing approach to begin identifying candidate mechanisms of *Xenopus* vocal evolution. *X. laevis* and *X. petersii* are ideal subjects for this approach because while they are relatively closely related, male advertisement calls are significantly longer in duration and period in *X. laevis* compared to *X. petersii* (23). RNA sequencing of the vocal premotor nucleus PB, revealed that approximately 25% (6,539 of 26,576) of genes were differentially expressed between male *X. laevis* and *X. petersii*. Because *Xenopus* vocalizations are produced by sexually dimorphic and temporally dynamic central pattern generators (30,32), we focus our discussion on candidate genes involved in steroid hormone signaling, neuromodulation, and neuronal membrane and synaptic properties.

### Hormone signaling

The *Xenopus* hindbrain vocal circuit was initially identified due to its expression of androgen receptors (33–35). Consistent with previous studies, we detected high androgen-receptor expression in PB of both *X. laevis* and *X. petersii*, which was not differentially expressed between species. Although other steroid hormones have not been implicated in male vocal behaviors, we detected transcripts for several additional steroid nuclear receptors, including estrogen, progesterone, glucocorticoid, and mineralocorticoid. While estrogen receptor expression had previously been indicated in *X. laevis* midbrain and forebrain in both sexes (36), it had not been described in PB. These results indicate a potentially complex system of steroid regulation of male vocal behaviors. While the role of androgens in development and maintenance of male *X. laevis* advertisement calling has been well-studied for decades (30,37–39), the involvement of other steroid hormones and their receptors has been largely overlooked and warrants careful examination.

Further supporting a central role for steroid hormones in vocal regulation, we identified many steroid synthesis and metabolic enzyme transcripts in both species. These results indicate that steroids may be synthesized locally in the vocal circuit, as observed in other species (40–43). Examples of relationships between steroidogenic enzymes and behaviors have been shown between species (in weakly electric fish; (44), reproductive morphs (in ruff sandpipers; (45), and behaviorally distinct individuals (*C. elegans*; (46)). Thus, the species differences in local steroid synthesis in the *Xenopus* PB provides a promising opportunity to explore the degree to which differential expression of steroid metabolizing enzymes can contribute to behavioral evolution.

In addition to steroid hormones, prostaglandins have been implicated in vocal communication across the frog phylogeny. For example, exogenous prostaglandin exposure inhibited mate calling (*Bufo americanus* and *Rana pipiens;* (47) and release calling (*X. laevis*, *R. catesbiana*, *R. pipiens;* (*48–50*), and promoted phonotaxis behaviors in (*B. americanus*, *Hyla chrysocelis*, and *H.versicolor*; (51–53). In addition, preliminary evidence suggested a direct action of prostaglandins in the *R. pipiens* PB (54; referred to as the pretrigeminal nucleus in that study). More broadly, prostaglandins have been implicated in reproductive behaviors in multiple teleost fish species (55–57). The finding that prostaglandin pathway genes were also prominent in the current dataset indicates a likely role for vocal regulation in male *Xenopus* vocal behaviors, providing another source of behavioral variability between *X. laevis* and *X. petersii*.

### Neuropeptide and neuromodulator signaling

#### Neuropeptides

Arginine vasopressin (AVP, also called arginine vasotocin in frogs) has a well-described role in regulation of frog vocal behaviors (58). In various frog species, exogenous AVP has been shown to induce changes in call type (59), call rate (58), and amount of time spent calling (60,61). Effects on vocal behaviors are found beyond frogs – for example in midshipman vocal fish, AVP application inhibits calling in courting males (62). Changes in AVP receptor expression drive behavioral differences between monogamous prairie voles and promiscuous montane voles (63), while altered expression of the AVP peptide between mouse species is known to affect nest building behaviors (64). Expression of AVP and its receptors in *X. laevis* and *X. petersii* indicates that vasopressin regulates calling in both species, while changes in AVP pathway genes could underlie vocal differences between species.

The link between oxytocin and social communication is strong across taxa, though evidence for direct action on vocal production circuits is limited. In *E. coqui* frogs, oxytocin injection was shown to increase call rate (60). In midshipman fish the teleost oxytocin homolog (isotocin) inhibits calling (62). In zebra finches, oxytocin-containing fibers have been observed in key vocal nuclei including HVC (65) and oxytocin antagonists can decrease singing (66). Previous work in *X. laevis* revealed oxytocin-immunoreactive fibers in the area of PB (67) The current finding of differentially expressed oxytocin receptors indicate a likely role for oxytocin in regulating species-typical *Xenopus* call patterns.

While galanin neurons have been shown to regulate parental behaviors in mammals (68), there is only limited evidence suggesting a role in vertebrate vocal production (69). Differential expression of galanin receptors in the current study indicates that galanin may act on the *Xenopus* vocal circuit, with potentially divergent effects in *X. laevis* and *X. petersii*. Substance P has been shown to modulate vocal neurons in *X. laevis* PB (70) and respiratory rhythms in a putative PB homolog in lamprey (71). In this study the genes encoding substance P and its receptor were highly, but not differentially, expressed, supporting a conserved role in both *X. laevis* and *X. petersii*.

#### Monoamine neuromodulators

Neuromodulators can have profound effects on circuit activity across a range of timescales (72,73), and changes in neuromodulator signaling can therefore drive behavioral evolution across species (5,74). Short-term behavioral effects of monoamine neuromodulators (e.g. serotonin, dopamine, and norepinephrine) have been demonstrated across species including zebra finches and midshipman fish (75–79). Monoamine-dependent behavioral changes during development have also been described, for example in *Xenopus* tadpoles in which increased swimming frequency correlates with the upregulation of spinal D1-like dopamine receptors (80). While direct evidence for monoamine involvement in driving vocal evolution is limited, evidence from other behavioral systems supports a potential role. In three-spine sticklebacks, for example, marine and freshwater populations exhibit altered expression of adrenergic, serotonergic, and dopaminergic receptors (81), providing possible mechanisms for behavioral differences such as tendency to school (82) and aggression (83). Thus monoamines and their receptors are good candidates for promoting species-typical behaviors during behavioral evolution.

The current study revealed the expression of serotonergic, adrenergic, and dopaminergic receptors in PB of both species (Fig. 4C). The vocal circuit in *Xenopus* can be activated *in vitro* with serotonin application, which initiates fictive calling via serotonin 2C receptors (5HT2C; (84,85). Confirming previous findings, 5HT2C genes were the most highly expressed serotonin receptor genes in both species and were not differentially expressed. However, 21 additional serotonin receptor genes were identified(including 5 DEGs). Thus, it is likely that serotonin plays additional modulatory roles through other receptor subtypes. Although dopaminergic and adrenergic signaling are not known to impact *Xenopus* vocalizations, the presence of differentially expressed dopaminergic and adrenergic receptor genes provides prime gene candidate targets for investigating whether these neuromodulator systems underlie species differences in *Xenopus* vocal pattern generation.

### Ion channels and neurotransmitter receptors

#### NMDA receptor currents

A population of PB neurons in both *X. laevis* and *X. petersii* exhibit NMDA receptor-dependent oscillations that are thought to underlie species-typical call timing (7). NMDAR-dependent oscillations have been identified in a variety of organisms and circuits, highlighting their importance in rhythm generation (86–95). We therefore hypothesized that species differences between neurons depend on distinct expression of NMDA receptors (NMDARs) and other ion channels that regulate the timing of these oscillations.

One possible mechanism for shorter membrane potential oscillations is *X. petersii* neurons could involve NMDAR subunits with higher calcium conductances, which could shorten the delay between NMDAR-dependent depolarization and activation of a calcium-dependent outward current leading to repolarization. Here, we found that two NMDAR genes (encoding GluN2D and GluN3B) are upregulated in *X. petersii*. GluN3B subunits (upregulated in *X. petersii)* have been shown to form triheteromeric complexes with elevated calcium permeabilities (96), providing a potential mechanism for shortened oscillations in *X. petersii*. Future physiology studies can test the hypothesis that *X. petersii* neurons exhibit increased NMDAR calcium conductances that in turn generate shortened membrane oscillations.

#### Inward currents

L-type calcium currents have been shown to regulate NMDAR-dependent oscillation duration (97). Here we found differential expression of voltage-gated gamma subunits known to modulate L-type calcium channel activation and inactivation (98), providing another possible mechanism of species-typical membrane potential oscillations.

Calcium-dependent nonselective cation currents (I_CAN_) can also contribute to rhythmic bursting (99–101). While I_CAN_ is thought to be primarily mediated by TRPM4 in mammalian respiratory circuits (102) and TRPM5 in mammalian locomotor circuits (103), other TRP channels may contribute to rhythmic pattern generation. For example, TRPC3 has been shown to contribute to pacemaker properties of dopaminergic substantia nigra neurons (104). While TRPM4 and TRPM5 are not expressed in the current dataset, eight other differentially expressed TRP channel genes were detected including trpc3.S, which is downregulated in *X. petersii* and could contribute to shorter depolarization durations.

Persistent sodium currents (I_NaP_; (99,105–110) and hyperpolarization-activated cation current (I_h_; (111–113) have also been shown to contribute to rhythm generation in respiratory and motor circuits. Ion channels underlying these currents were present in *X. petersii* and *X. laevis,* supporting their involvement in vocal pattern generation; however they were not differentially expressed and are unlikely to drive species differences.

#### Outward currents

Genes underlying outward currents shown to underlie rhythmic oscillations and bursting are also expressed in PB. Of particular interest are the calcium-dependent potassium currents, which are important for driving repolarization (114–120). The large-conductance calcium-activated potassium channel (BK) gene (kcnma1.L) was highly (but not differentially) expressed, indicating an important role for BK in both species. However, two of the five genes coding for BK 𝛽 subunits were differentially expressed: kcnmb3.S (encoding 𝛽3) was upregulated, and kcnmb1.S (encoding 𝛽1) was downregulated, in *X. petersii*. Interestingly, human BK channel α subunits co-expressed with the 𝛽3 subunit have faster activation and inactivation kinetics than when co-expressed with 𝛽1 subunits (121). In another study, the 𝛽3 subunit caused a rapidly inactivating current while 𝛽1 co-expression (upregulated in *X. laevis*) showed little inactivation (122) Thus, the observed 𝛽1 and 𝛽3 expression differences could collectively support species-typical oscillation kinetics (faster in *X. petersii*, slower in *X. laevis*).

Other outward currents may regulate temporal patterns of rhythmic oscillations, including A-type current (Kv4 and Kv1; (123), M-type current (Kv7; (124–126), and delayed rectifier currents (Kv2.1 and 2.2 and silent modifiers Kv9.1 and Kv9.3; (127). The delayed rectifier silent modifier Kv9.3 channel subunit gene (kcns3.S) – upregulated in *X. petersii* – can form heteromeric Kv2.1 channels that lead to faster recovery from inactivation (128); increased expression in *X. petersii* could therefore support a briefer call period compared to *X. laevis.* Two genes underlying the Kv7-dependent M current were upregulated in *X. petersii*. In rodents, blocking Kv7 channels lengthens neuronal burst duration and period in locomotor CPG interneurons and slows locomotion (124–126), suggesting that increased expression of Kv7 channels in *X. petersii* could promote shorter trill duration and period via enhanced M current.

#### Transmitter receptors

In addition to NMDARs, we found differential expression of excitatory and inhibitory neurotransmitter receptors, including glutamate, nicotinic acetylcholine, and GABA receptors. These candidate genes serve as strong candidates for testing the degree to which evolution of synaptic features shapes species differences in behavior (e.g. 1,2).

### Summary

Our study revealed hundreds of differentially expressed genes that represent prime candidates for underlying species differences in *X. laevis* and *X. petersii* call timing. These results support new hypotheses regarding the mechanisms of vocal production and evolution in *Xenopus* and across taxa. While the current candidate genes arise from bulk RNA sequencing data, hypotheses about the role of these genes can be tested through established behavioral, physiological, and molecular profiling approaches at the level of whole organisms, circuits, and single neurons. However, changes in gene expression are likely only one of several mechanisms of behavioral evolution (74,129,130). In particular, given the relatively recent genome duplication in the ancestors of *X. laevis* and *X. petersi*i approximately 17 million years ago (131), neofunctionalization is another likely process driving *Xenopus* vocal evolution as has been demonstrated in other taxa (17,132,133). Given the ability to integrate established behavioral and physiological approaches with increasingly powerful molecular profiling technologies, the *Xenopus* vocal system offers a promising avenue for identifying common mechanisms underlying the evolution of behavioral circuits.

## SUPPLEMENTAL MATERIALS

**Supplemental table 2.1.**

| SampleName | InputReads | UnqMap | UnqMapPerc | MultiMap | MultiMapPerc | TooManyLoci | TooManyLociPerc | UnmappedTooManyMismatch | UnmappedTooShort | UnmappedOther |
| --- | --- | --- | --- | --- | --- | --- | --- | --- | --- | --- |
| XL1 | 95,440,158 | 87,489,968 | 91.67% | 2,778,172 | 2.91% | 335,527 | 0.35% | 0.00% | 4.64% | 0.43% |
| XL10 | 94,415,256 | 86,521,153 | 91.64% | 2,965,869 | 3.14% | 350,337 | 0.37% | 0.00% | 4.37% | 0.47% |
| XL11 | 82,695,181 | 76,189,874 | 92.13% | 2,393,785 | 2.89% | 291,097 | 0.35% | 0.00% | 4.18% | 0.44% |
| XL12 | 97,808,538 | 89,796,296 | 91.81% | 2,920,149 | 2.99% | 356,476 | 0.36% | 0.00% | 4.38% | 0.46% |
| XL13 | 102,253,093 | 93,685,561 | 91.62% | 3,068,809 | 3.00% | 382,077 | 0.37% | 0.00% | 4.51% | 0.50% |
| XL4 | 88,735,031 | 81,337,915 | 91.66% | 2,732,612 | 3.08% | 328,086 | 0.37% | 0.00% | 4.44% | 0.45% |
| XL5 | 110,323,566 | 100,775,326 | 91.35% | 3,533,361 | 3.20% | 429,934 | 0.39% | 0.00% | 4.57% | 0.49% |
| XL6 | 75,790,621 | 69,081,499 | 91.15% | 2,397,319 | 3.16% | 302,006 | 0.40% | 0.00% | 4.79% | 0.50% |
| XL7 | 87,404,388 | 80,111,156 | 91.66% | 2,672,826 | 3.06% | 325,725 | 0.37% | 0.00% | 4.45% | 0.47% |
| XL8 | 89,215,499 | 81,541,571 | 91.40% | 2,802,298 | 3.14% | 347,327 | 0.39% | 0.00% | 4.56% | 0.51% |
| XP1 | 99,658,388 | 83,444,037 | 83.73% | 4,693,483 | 4.71% | 547,890 | 0.55% | 0.00% | 10.42% | 0.60% |
| XP10 | 89,904,906 | 76,843,407 | 85.47% | 3,957,319 | 4.40% | 423,826 | 0.47% | 0.00% | 9.14% | 0.52% |
| XP11 | 66,149,856 | 56,675,193 | 85.68% | 2,835,360 | 4.29% | 312,675 | 0.47% | 0.00% | 9.07% | 0.49% |
| XP13 | 78,734,039 | 67,628,325 | 85.89% | 3,322,886 | 4.22% | 328,212 | 0.42% | 0.00% | 9.00% | 0.47% |
| XP3 | 88,295,645 | 74,404,277 | 84.27% | 4,084,950 | 4.63% | 451,254 | 0.51% | 0.00% | 10.03% | 0.56% |
| XP4 | 106,351,986 | 89,225,740 | 83.90% | 4,939,847 | 4.64% | 581,291 | 0.55% | 0.00% | 10.34% | 0.57% |
| XP5 | 86,735,701 | 73,092,619 | 84.27% | 4,039,949 | 4.66% | 487,786 | 0.56% | 0.00% | 9.94% | 0.57% |
| XP7 | 99,831,315 | 85,397,378 | 85.54% | 4,390,565 | 4.40% | 496,198 | 0.50% | 0.00% | 9.06% | 0.50% |
| XP8 | 79,042,847 | 67,566,723 | 85.48% | 3,452,693 | 4.37% | 364,959 | 0.46% | 0.00% | 9.22% | 0.47% |
| XP9 | 72,699,049 | 62,090,070 | 85.41% | 3,178,041 | 4.37% | 333,630 | 0.46% | 0.00% | 9.28% | 0.48% |

**Supplemental data table 2.1.**

| ID | Go Term Description | Adjusted p value | Count |
| --- | --- | --- | --- |
| GO:0004497 | monooxygenase activity | 0.000001 | 42 |
| GO:0005506 | iron ion binding | 0.000001 | 54 |
| GO:0020037 | heme binding | 0.000007 | 48 |
| GO:0046906 | tetrapyrrole binding | 0.000028 | 48 |
| GO:0005102 | signaling receptor binding | 0.000202 | 117 |
| GO:0016705 | oxidoreductase activity | 0.000274 | 51 |
| GO:0004175 | endopeptidase activity | 0.003891 | 91 |
| GO:0016701 | oxidoreductase activity | 0.005923 | 11 |
| GO:0016702 | oxidoreductase activity | 0.005923 | 11 |
| GO:0004867 | serine-type endopeptidase inhibitor activity | 0.009671 | 19 |
| GO:0004364 | glutathione transferase activity | 0.010396 | 9 |
| GO:0005179 | hormone activity | 0.011467 | 28 |
| GO:0048018 | receptor ligand activity | 0.012187 | 64 |
| GO:0030546 | signaling receptor activator activity | 0.013098 | 64 |
| GO:0030545 | signaling receptor regulator activity | 0.013772 | 65 |
| GO:0004930 | G protein-coupled receptor activity | 0.013772 | 126 |
| GO:0004222 | metalloendopeptidase activity | 0.017532 | 41 |
| GO:0005523 | tropomyosin binding | 0.032181 | 7 |
| GO:0005201 | extracellular matrix structural constituent | 0.035412 | 13 |
| GO:0016712 | oxidoreductase activity | 0.054317 | 10 |
| GO:0099094 | ligand-gated monoatomic cation channel activity | 0.057913 | 20 |
| GO:0004953 | icosanoid receptor activity | 0.059672 | 7 |
| GO:0008237 | metallopeptidase activity | 0.099137 | 58 |
| GO:0061134 | peptidase regulator activity | 0.099137 | 27 |
| GO:0005126 | cytokine receptor binding | 0.099137 | 18 |
| GO:0005164 | tumor necrosis factor receptor binding | 0.099137 | 10 |
| GO:0032813 | tumor necrosis factor receptor superfamily binding | 0.099137 | 10 |
| GO:0140375 | immune receptor activity | 0.099137 | 16 |
| GO:0003887 | DNA-directed DNA polymerase activity | 0.099137 | 9 |
| GO:0030414 | peptidase inhibitor activity | 0.099137 | 26 |
| GO:0051213 | dioxygenase activity | 0.099137 | 27 |
| GO:0004866 | endopeptidase inhibitor activity | 0.104512 | 24 |
| GO:0001664 | G protein-coupled receptor binding | 0.111713 | 16 |
| GO:0061135 | endopeptidase regulator activity | 0.119651 | 24 |
| GO:0090482 | vitamin transmembrane transporter activity | 0.119662 | 6 |
| GO:0034061 | DNA polymerase activity | 0.124919 | 9 |
| GO:0005540 | hyaluronic acid binding | 0.125616 | 7 |
| GO:0015267 | channel activity | 0.131532 | 86 |
| GO:0022803 | passive transmembrane transporter activity | 0.131532 | 86 |
| GO:0005539 | glycosaminoglycan binding | 0.141104 | 18 |

**Supplemental data table 2.2.**

| ID | Go Term Description | Adjusted p value | Count |
| --- | --- | --- | --- |
| GO:0002376 | immune system process | 0.000163 | 70 |
| GO:0006955 | immune response | 0.000163 | 55 |
| GO:0006952 | defense response | 0.006036 | 44 |
| GO:0002253 | activation of immune response | 0.006036 | 20 |
| GO:0098542 | defense response to other organism | 0.006036 | 34 |
| GO:0009607 | response to biotic stimulus | 0.010782 | 35 |
| GO:0002764 | immune response-regulating signaling pathway | 0.010782 | 15 |
| GO:0043207 | response to external biotic stimulus | 0.010782 | 34 |
| GO:0044419 | biological process involved in interspecies interaction between organisms | 0.010782 | 34 |
| GO:0051707 | response to other organism | 0.010782 | 34 |
| GO:0045087 | innate immune response | 0.011158 | 28 |
| GO:0002757 | immune response-activating signaling pathway | 0.012808 | 14 |
| GO:0050778 | positive regulation of immune response | 0.014977 | 20 |
| GO:0050776 | regulation of immune response | 0.035697 | 22 |
| GO:0009605 | response to external stimulus | 0.039358 | 57 |
| GO:0003008 | system process | 0.039358 | 44 |
| GO:0051694 | pointed-end actin filament capping | 0.048704 | 7 |
| GO:0002682 | regulation of immune system process | 0.049949 | 28 |
| GO:0002684 | positive regulation of immune system process | 0.094984 | 21 |
| GO:0002224 | toll-like receptor signaling pathway | 0.111206 | 10 |
| GO:0007051 | spindle organization | 0.119347 | 18 |
| GO:0019751 | polyol metabolic process | 0.128893 | 20 |

**Supplemental data table 2.3.**

| ID | Go Term Description | Adjusted p value | Count |
| --- | --- | --- | --- |
| GO:0005581 | collagen trimer | 0.004295 | 27 |
| GO:0030312 | external encapsulating structure | 0.023254 | 31 |
| GO:0031012 | extracellular matrix | 0.023254 | 31 |
| GO:0005615 | extracellular space | 0.023254 | 47 |

**Supplemental data table 3.**

| ID | KEGG Term Description | Adjusted p value | Count |
| --- | --- | --- | --- |
| xla00982 | Drug metabolism | 0.000004 | 22 |
| xla04820 | Cytoskeleton in muscle cells | 0.000004 | 112 |
| xla00140 | Steroid hormone biosynthesis | 0.000032 | 26 |
| xla04512 | ECM | 0.000169 | 46 |
| xla00980 | Metabolism of xenobiotics by cytochrome P450 | 0.000294 | 21 |
| xla04060 | Cytokine | 0.000441 | 68 |
| xla00983 | Drug metabolism | 0.001571 | 29 |
| xla00830 | Retinol metabolism | 0.005988 | 20 |
| xla00480 | Glutathione metabolism | 0.007652 | 28 |
| xla00590 | Arachidonic acid metabolism | 0.025187 | 22 |
| xla04672 | Intestinal immune network for IgA production | 0.048596 | 12 |
| xla01232 | Nucleotide metabolism | 0.051533 | 38 |
| xla00230 | Purine metabolism | 0.060823 | 56 |
| xla00760 | Nicotinate and nicotinamide metabolism | 0.074333 | 19 |
| xla00040 | Pentose and glucuronate interconversions | 0.076653 | 11 |
| xla00430 | Taurine and hypotaurine metabolism | 0.076653 | 11 |
| xla00250 | Alanine, aspartate and glutamate metabolism | 0.096486 | 19 |
| xla00350 | Tyrosine metabolism | 0.117504 | 14 |
| xla00910 | Nitrogen metabolism | 0.119249 | 8 |
| xla04622 | RIG | 0.130396 | 30 |
| xla04270 | Vascular smooth muscle contraction | 0.134999 | 56 |
| xla00790 | Folate biosynthesis | 0.182816 | 13 |

